# A Network-level Test of the Role of the Co-activated Default Mode Network in Episodic Recall and Social Cognition

**DOI:** 10.1101/2021.01.08.425921

**Authors:** Rebecca L. Jackson, Gina F. Humphreys, Grace E. Rice, Richard J. Binney, Matthew A. Lambon Ralph

## Abstract

Resting-state network research is extremely influential, yet the functions of many networks remain unknown. In part, this is due to typical (e.g., univariate) analyses testing the function of individual regions and not the full set of co-activated regions that form a network. Connectivity is dynamic and the function of a region may change based on its current connections. Therefore, determining the function of a network requires assessment at the network-level. Yet popular theories implicating the default mode network (DMN) in episodic memory and social cognition, rest principally upon analyses performed at the level of individual brain regions. Here we use independent component analysis to formally test the role of the DMN in episodic and social processing at the network level. As well as an episodic retrieval task, two independent datasets were employed to assess DMN function across the breadth of social cognition; a person knowledge judgement and a theory of mind task. Each task dataset was separated into networks of co-activated regions. In each, the co-activated DMN, was identified through comparison to an *a priori* template and its relation to the task model assessed. This co-activated DMN did not show greater activity in episodic or social tasks than high-level baseline conditions. Thus, no evidence was found to support hypotheses that the co-activated DMN is involved in explicit episodic or social processing tasks at a network-level. The networks associated with these processes are described. Implications for prior univariate findings and the functional significance of the co-activated DMN are considered.

## Introduction

Most higher cognitive functions are not localised to a single brain region but rather reflect the joint action of a distributed network of regions (Mesulam 1990; Catani et al. 2012; Humphreys et al. 2014). As functional connectivity is dynamic, individual regions may form part of different networks across time (Breakspear 2004; Hutchison et al. 2013). The role of a brain area is determined by its current ‘neural context’; its interactions with other connected regions (McIntosh 1999; Chen et al. 2017). Thus, the relationship between a single brain region and a function may change across time as it is embedded within different networks (Hagoort 2014). Therefore, it is imperative to understand the functional significance of brain networks by assessing their relationship to functions at the network-level (and not only test associations between functions and individual brain regions; Smith et al. 2009; Laird et al. 2011; Leech et al. 2011; Geranmayeh et al. 2012; Geranmayeh et al. 2014; Hagoort 2014; Jackson et al. 2019).

Networks of co-activated brain regions can be extracted from the time-course of resting-state fMRI data (i.e., data acquired without a formal task) using Independent Component Analysis (ICA; Beckmann et al. 2005; Calhoun et al. 2008; Smith et al. 2009; Geranmayeh et al. 2012; Geranmayeh et al. 2014). The ease of collection of resting-state fMRI potentially licenses the use of these ‘resting-state networks’ (RSNs) in both basic and clinical neurosciences and has led to a recent explosion of interest in studying networks across different disciplines (Rosazza et al. 2011; de la Iglesia-Vaya et al. 2013). This has greatly improved our understanding of network organisation in the brain, and is potentially informative about network-function relationships as RSNs reflect connectivity patterns identifiable across multiple task states (earning them the alternative name ‘Intrinsic Connectivity Networks’; Smith et al. 2009; Laird et al. 2011). However, resting-state analyses are inherently detached from function due to the lack of functional information during resting-state scans, which has slowed the association of these networks with cognitive functions. Theories of RSN function are typically based on comparison to task data. Yet, despite the interest in understanding the function associated with networks, most task-based analyses rely on methods which assess the function of regions not networks. This results in a disconnection between the methods used to relate functions and the brain, which are typically at the region-level (including univariate analyses and meta-analyses), and the assertions made, which are often at the network-level. As the role of constituent regions change over time, making assertions about network-function relationships requires network-level approaches to understand function (Smith et al. 2009; Laird et al. 2011; Leech et al. 2011; Geranmayeh et al. 2012; Geranmayeh et al. 2014; Hagoort 2014; Jackson et al. 2019), for instance, detecting co-activated sets of regions in task data with ICA and examining their relation to the task. Identifying components that reflect known RSNs and probing their function can provide a powerful test of existing hypotheses of the function of a network, generated from analyses of the constituent regions.

There is great interest in the possible function of one RSN in particular, the default mode network (DMN). The DMN was initially defined as the areas that were more active during rest than in most tasks (Shulman et al. 1997; Raichle et al. 2001), yet has been shown to reflect a network of co-activated regions that can be identified during rest and task states (Greicius et al. 2003; Buckner et al. 2008). Areas associated with the DMN include medial prefrontal cortex, posterior cingulate cortices, precuneus, angular gyrus and lateral and medial temporal cortices, although the central focus is the medial surface (Raichle et al. 2001; Buckner et al. 2008; Utevsky et al. 2014). The term ‘default mode network’ is typically used to refer to both the set of regions which often deactivate during many tasks and the network of co-activated regions which are engaged together. Whilst there is great overlap between the areas implied by these two meanings, it is important to distinguish between them to make assertions at the appropriate (network or region) level for a given analysis. There is clear evidence for the functional roles of many of the relevant regions, yet there have been relatively few direct tests of the function of the joint activity of the co-activated network, here referred to as the ‘co-activated DMN’, which is the focus of the current investigation. Furthermore, these definitions may lead to some differences in spatial extent as, whilst there is a consistent core of regions which researchers agree are part of the co-activated DMN, there is evidence that some task-negative regions belong to distinct networks (e.g., the anterior temporal lobes; Humphreys et al. 2015; Jackson et al. 2019).

The function of the DMN is highly debated. Finding greater activity in DMN regions in rest compared to various tasks led to the theory that the DMN is simply performing some process that occurs frequently during free thought, yet would occur in an active task if the appropriate domain was tested (Buckner et al. 2008; Callard et al. 2012; Spreng 2012). Within this framework three popular candidate processes have been postulated; episodic memory, social cognition and semantic cognition (Binder et al. 1999; Buckner et al. 2008; Binder et al. 2009; Mars et al. 2012). Evidence in favour of a critical role of the DMN in these processes was predominantly gained from region-level assessments, which generate a map of the areas involved in a process based on univariate or meta-analyses, which is then compared to the extent of the DMN (often through simple visual comparison; e.g., Binder et al. 1999; Buckner et al. 2008; Binder et al. 2009; Kim 2010; Mars et al. 2012; DiNicola et al. 2020). These analyses do not directly assess the function of the network and are prone to various sources of error due to the additional steps required to move from region-level to network-level assessments. For instance, when comparing the extent of the network and the task-based result, there is a question of how to judge quantitatively whether spatial maps of a network and a univariate analysis result are sufficiently similar to reflect the same network. However, a much greater concern is that differences in task activation may occur in regions overlapping those in a network without involvement of the co-activated network in that domain.

As regions may be involved in different networks responsible for different functions (Breakspear 2004; Hutchison et al. 2013), the involvement of one or more areas in a task does not necessitate the importance of a specific associated network. A region may be involved in a given function only when associated with one particular set of connections, which may or may not be the network of interest. Additionally, an activation difference within one region could be due to the summed activity of two networks that overlap in that area. Region-level methods cannot distinguish the function of multiple, divisible RSNs, instead they merely average over the different functional profiles a region may have whilst involved in different networks. An additional problem that may affect the DMN in particular, is that task difficulty differences can result in the identification of areas responsible for task-general processing or that show task general deactivation, typically thought to involve similar areas to known resting-state networks (Vincent et al. 2008; Duncan 2010; Gilbert et al. 2012; Humphreys et al. 2014; Assem et al. 2020; Humphreys et al. in press). Furthermore, if a task-related process is performed quicker in one condition, the extra remaining time within a trial means a greater amount of rest or off-task processing might occur. This means that RSNs incidental to the task (particularly the DMN which is preferentially involved in rest) may show greater activity in this condition. This could explain the broad similarity between the DMN and the results of meta-analyses of episodic memory, social cognition and semantic cognition (Binder et al. 1999; Buckner et al. 2008; Binder et al. 2009; Mars et al. 2012) without an active role for the DMN in these processes, if there are systematic differences in the difficulty of the contrasts typically employed (Humphreys et al. 2017; Humphreys et al. in press). Partial-least squares analyses demonstrate involvement of sets of areas overlapping the DMN in episodic and social cognition, and have some advantages over univariate analyses as they identify covariation across the brain (Spreng et al. 2010a; Spreng et al. 2010b; Hassabis et al. 2014; DuPre et al. 2016). However, they remain subject to many of these concerns (e.g., combining multiple networks) due to the identification of a single set of regions whose activation differs between two task conditions.

The difficulty associating the co-activated DMN with a particular function on the basis of region-level assessments has been demonstrated in the domain of semantic cognition (Jackson et al. 2019). There is great overlap between the DMN and the results of a large meta-analysis of semantics, resulting in the popular hypothesis that the DMN is recruited for semantic cognition (Binder et al. 2009; Binder et al. 2011), albeit with some authors highlighting regional differences in responsiveness to semantic and non-semantic tasks (e.g., Humphreys et al. 2017). However, a recent study using ICA to identify the co-activated DMN, did not find results consistent with this proposed role in semantic cognition. Instead, the co-activated DMN demonstrated similar activation levels for semantic and non-semantic tasks, with both deactivated compared to rest. A distinct ‘semantic network’ showed greater activation for a semantic than non-semantic task (Jackson et al. 2019). In the present study, we used the same network-level approach to directly evaluate the hypotheses that the co-activated DMN is responsible either for episodic or social processing. This network-level approach entails 1) using ICA to detect the different sets of regions that co-activate in relevant task data, 2) using quantitative overlap measures with *a priori* DMN templates to identify which set of regions reflects the co-activated DMN and 3) examining the relationship between the time course of this co-activated DMN and the relevant task contrast. As social cognition may be multi-factorial (Van Overwalle 2009; Mars et al. 2012; Rice et al. 2018), two tasks evaluating diverse aspects of social cognition were employed; one testing the semantic processing of person knowledge and one engaging theory of mind processes.

## Methods

Three previously-reported fMRI datasets including episodic and social tasks were analysed. These included one *Episodic Task* (Humphreys et al. in press), one *Social - Person Knowledge Task* (Rice et al. 2018) and one *Social - Theory of Mind Task* (Barch et al. 2013; Van Essen et al. 2013). The *Episodic* and the *Social - Person Knowledge* datasets employed dual echo EPI fMRI acquisition, improving signal in the problematic inferior frontal and temporal regions (including key medial prefrontal cortex areas) without loss of signal elsewhere (Poser et al. 2006; Halai et al. 2014). The *Social - Theory of Mind Task* was downloaded from the Human Connectome Project (HCP; Barch et al. 2013; Van Essen et al. 2013). This does not have the benefit of dual echo EPI fMRI, but has a large number of participants, a high spatial resolution and processing steps designed to correct for signal distortion (Glasser et al. 2013; Van Essen et al. 2013).

The multi-step procedure outlined in Jackson et al., (2019) was followed. Analyses were performed separately for each task dataset. First, ICA was performed to determine the networks present within the data. Secondly, the resting-state network-of-interest was identified through quantitative comparison of each resulting component to a template. This avoids the researcher bias present in visual identification and helps determine whether one or more task components relates to the network-of-interest, i.e., whether the network has split and formed multiple components. In the final step, the functional role of the network-of-interest was tested by comparison to the task model. This provides a test of the function of the network *per se* and not merely the constituent regions, as for instance, univariate analyses would. For each task, a single contrast was used to determine whether the network-of-interest was involved in the condition of interest. These critical contrasts compared the task condition of interest to an active baseline as comparison to rest may be affected by the propensity of the process to occur during rest (Frackowiak 1991; Visser et al. 2010; Jackson et al. 2019). In particular, the DMN is expected to be prevalent during rest, yet if it is responsible for episodic or social processing it should also be prevalent during these tasks to a greater extent than high-level control tasks. However, all plots are shown against rest for ease of interpretability across tasks.

### Participants

In the *Episodic Task* data, 22 participants completed the study (18 females; mean age 24.55, SD 4.74). All were included in the present analyses. In the *Social - Person Knowledge Task*, data were acquired for 20 participants (16 female; mean age = 26.6 years, range = 20-42). One participant was excluded due to excessive motion (see Preprocessing Methods). The participant groups were independent with no participants involved in both studies. All had normal or corrected-to-normal vision and hearing, and no history of psychiatric or neurological disorder. Both studies were approved by the local ethics board and all participants provided written informed consent.

The *Social - Theory of Mind Task* data was a subset of 80 participants of the Human Connectome Project 1200 Subject release (Van Essen et al. 2013). This number far exceeds the number of participants typically used in an fMRI study. Inclusion criteria for the HCP consisted of an age between 22 and 35 years and no significant history of psychiatric disorder, substance abuse, neurological, or cardiovascular disease. The first 40 male and 40 female participants with no data collection or analysis quality control issues were included in this study. All had an acquisition site of Washington University.

### Imaging protocol

The *Episodic* and *Social - Person Knowledge* studies used the same scanner and a very similar acquisition protocol. Participants were scanned using a Phillips Achieva 3.0T system with a 32 channel SENSE coil with a sense factor of 2.5 whilst wearing noise cancelling Mk II+ headphones (MR Confon, Magdeburg, Germany). Structural references were obtained with an in-plane resolution of .938 and slice thickness of 1.173mm. Both studies utilised a dual gradient echo EPI technique to reduce signal dropout in inferior temporal and frontal cortex whilst maintaining contrast sensitivity throughout the brain (Poser et al. 2006; Poser et al. 2007, 2009; Halai et al. 2014). This involved parallel acquisition at a short (12ms) and a long (35 ms) echo time, and linear summation of the resulting images. The whole brain was covered (field of view 240x240mm, resolution matrix 80x80 for the *Social - Person Knowledge Task* and 80x79 for the *Episodic Task*, reconstructed voxel size of 3mm and slice thickness of 4mm) with a tilt of 45° off the AC-PC line to reduce the effect of ghosting in the temporal lobe. The flip angle was 85° and the TR was 2.8.

The *Social -Theory of Mind* dataset was acquired using the HCP stage 2 protocol, described in full in Uğurbil et al., (2013). In brief, the use of highly-accelerated, multiplexed, multi-banding EPI (Feinberg et al. 2010; Moeller et al. 2010; Setsompop et al. 2012; Xu et al. 2012) allowed a high spatial resolution (2.0 mm isotropic voxels) with a low TR (720 ms). The whole-brain image was acquired on a modified 3T Siemens Skyra with a 32 channel head coil (multi-band acceleration factor = 8, TE = 33.1 ms, flip angle = 52°, field of view = 208 × 180 mm, 72 slices). One run was acquired with a right-to-left phase encoding and the other with a left-to-right phase encoding, to help factor out the effect of phase encoding direction.

### Episodic Task Data

The *Episodic Task* data were previously reported in a univariate assessment of the role of inferior parietal regions in episodic and semantic memory (Humphreys et al. in press). The dataset was event-related and included episodic retrieval, semantic retrieval, semantic decision and visual (scrambled pattern decision) control conditions. The episodic retrieval task involved retrieving specific features of photographs presented prior to entering the scanner. Outside of the scanner participants were presented with a colour photograph of an object or scene for ten seconds and required to describe it aloud. Within the scanner, a question was presented referring to the previously seen object, for instance, ‘What colour was the balloon?’. Two potential answers were presented alongside the question (e.g., ‘Blue’ and ‘Green’). A scrambled photo was also present on screen, but was not relevant to the performance of the episodic task. Participants reported the vividness of their memory on each trial on a scale from 1 to 4. Each episodic trial lasted 4 seconds. The visual control task involved a judgement of whether a scrambled photograph was on the left or right of the screen. There were three task runs each lasting 621.6 seconds and including 222 volumes. The task model included episodic, semantic retrieval, semantic decision, visual control, rest and instruction events. However, only the episodic and control conditions are of critical interest here as only these conditions attest to the proposed role of the DMN in episodic processing.

### Social - Person Knowledge Task Data

The data used to assess the role of the DMN in person knowledge were reported previously (‘Study 1’ in Rice et al. (2018)). In the original study, only univariate analyses were performed and the focus was the selectivity of social concepts (including multimodal person knowledge) within the anterior temporal lobes and not to probe the DMN. Participants performed the same semantic judgement (‘Is the stimuli European/not?’) on 3 different categories of stimuli (people, landmarks and animals) presented in either the auditory domain (as spoken words) or the visual domain (as pictures). Visual and auditory control conditions were created by scrambling the images and phase-scrambling auditory tones. Participants were asked whether the image was high or low on the screen or if the auditory tone was high or low-pitched. The *Social - Person Knowledge* task data were acquired over three 12 minute runs, each including 255 dynamic scans (including 2 dummy scans to be removed). Stimuli were presented in blocks of 18 seconds. Social (faces and names), non-social (auditory or visual presentation of landmarks and animals), control (visual and auditory baselines) and rest conditions were included in the task model. All contrasts included equal numbers of trials presented in the auditory and visual domains.

### Social - Theory of Mind Task Data

The *Social - Theory of Mind* data were downloaded from the HCP website (Van Essen et al. 2013). Full details of the task design are provided in Barch et al., (2013). Frith-Happé animations were taken from previous studies (Castelli et al. 2000; Wheatley et al. 2007) and, where necessary, shortened to a 20 second duration. These animations consist of moving shapes, which either move randomly or appear to interact with each other in a way that takes into account the other shapes thoughts and feelings. On each trial, participants viewed a social or random motion video for 20 seconds and were required to assess whether the shapes had an interaction of a social nature. Participants then responded that the shapes had a social interaction, did not have an interaction, or that they were not sure, in an additional 3 second window. This 23 second duration was treated as one block and 5 task blocks (mixed between social and random conditions) were presented with 5 fixation blocks in each of the 2 runs. Each run lasted 3 minutes and 27 seconds (including an 8 second task initiation period) and contained 274 volumes. The task model included social and non-social conditions, as well as rest. This task was chosen for the HCP as it gives reliable and robust activations across participants in univariate analyses and is considered an objective test of theory of mind processes (Castelli et al. 2000; Castelli et al. 2002; Wheatley et al. 2007; White et al. 2011; Barch et al. 2013). This is therefore an exemplary dataset in which to assess the role of the DMN in theory of mind processes.

### Preprocessing Methods

The *Episodic* and *Social - Person Knowledge* data were processed using a pipeline based on Jackson et al., (2019), involving the use of statistical parametric mapping (SPM 8) software (Wellcome Trust Center for Neuroimaging) and the Data Processing Assistant for Resting State fMRI (DPARSF Advanced Edition, V2.3) toolbox (Chao-Gan et al. 2010). This pipeline is based on resting-state processing and is designed to remove a substantial proportion of the effects of motion. However, unlike the resting-state processing, the global mean was not removed from the data and the data were not filtered, as these steps may change the relation between the activity of an area and the task model. The steps performed included slice time correction, realignment, coregistration, removal of motion and tissue-based regressors, normalisation using DARTEL and linear detrending. The tissue-based regressors included the average activity in the cerebrospinal fluid and the white matter. The motion regressors employed were the Volterra- expanded 24 motion parameters (Friston et al. 1996) and high motion time points identified using the ARtifact detection Tools software package (ART; www.nitrc.org/projects/artifact_detect). High motion trials, those with more than 1mm relative movement or where the mean intensity exceeded a z-score of 2.5, were removed. If removal of high motion trials resulted in the exclusion of more than a quarter of the remaining data then the participant was removed. This resulted in the removal of a single participant in the social dataset only. This motion correction procedure is more stringent than typical for task data preprocessing, but is typical for resting-state preprocessing due to the strong impact of motion on connectivity/co-activation measures and therefore is appropriate here.

The pre-processed *Social - Theory of Mind* data were downloaded from the HCP website (http://www.humanconnectomeproject.org/data/; Van Essen et al. 2013). The HCP processing pipeline is described in more detail in Glasser et al., (2013). The pipeline relies upon tools from FSL (Jenkinson et al. 2012) and FreeSurfer (Fischl 2012). The key steps are gradient unwarping, FLIRT-based motion correction, TOPUP-based distortion correction using the fieldmap, brain-boundary-based registration of the EPI data to the structural scan, non-linear (FNIRT) registration into MNI152 space, grand-mean intensity normalization and bias field removal. This process requires skull-stripping and segmentation of the structural scan using FreeSurfer. All data had to pass quality control checks to be included in the HCP dataset (see Marcus et al. 2013) and only those without issues identified were included in the present study. The resulting data are therefore, fully-processed high quality EPI data in MNI atlas space with care taken to minimise the effects of participant movement and distortion. As no overt smoothing had been performed on the downloaded images, smoothing was performed using a 4 mm full-width half maximum (FWHM) Gaussian kernel, following Barch et al., (2013).

### Independent Component Analyses

In order to determine the networks present, a separate group ICA was performed on each task dataset (including the whole scan and all runs). ICA was performed using the Group ICA of fMRI Toolbox (GIFT; Calhoun et al. 2001). The number of components within the data was estimated within GIFT using the group mean Minimum Description Length criteria (Li et al. 2006). The ICA was set to discover this number of components, deemed appropriate for the dataset. Each component comprises a spatial map and a time-course of activation. The resulting components are presented as group-level spatial maps, calculated by performing a one sample t-test on all the individuals’ data to identify voxels that are significantly involved across individuals. This may be performed using the utilities provided within GIFT and results in a standard statistical map for each component which was thresholded at a voxel-level significance threshold of .001 and an FWE-corrected critical cluster level of .05. Components are binarised (for the purposes of visualisation and template matching only) based on reaching significance at this threshold. Artefactual components were identified visually based on the spatial map of each component in the same manner as Jackson et al., (2019) and in line with published guidelines (Griffanti et al. 2017). This entailed removal of components with activity focussed outside of the grey matter (whether it be within the ventricles, white matter or outside the brain) or in the brainstem. When removing artefactual components, the researcher was blind to the correspondence of the components with the network templates and the task model. These artefactual components were excluded from further analyses.

### Identifying the Co-activated Networks in the Task Data

To ensure an accurate reflection of the consensus in the literature, two *a priori* templates were used to determine the component or components corresponding to the DMN in each task dataset; the DMN identified in the resting-state dataset in Jackson et al., (2019) and a publicly available DMN template (http://findlab.stanford.edu/functional_ROIs.html; Shirer et al. 2012). The two templates were used independently to identify the DMN and their results compared.

Identification of the DMN followed the same procedure as in Jackson et al., (2019). The Jaccard similarity coefficient (Jaccard 1912) was computed between a binarised map of the areas significantly active in the component at the group level and the binary templates. This gives a single measure of template fit that accounts for both extra and missing areas compared to the template, ranging from 0 (no overlap of active voxels) to 1 (full overlap of both active and inactive voxels). As in Jackson et al., (2019), a two-step procedure was employed, firstly allowing all components that reach a threshold of J = 0.15 to be identified and then assessing whether combining multiple components numerically improves the template fit. Unlike the sole use of a threshold, this two-step process allows identification of all components that may be related to the template and then clarifies whether multiple components are the best fit for the network (e.g., if the network has split) or if a single component is a better fit (e.g., there is only one DMN component and a distinct but spatially overlapping network). Note, Jaccard similarity scores are typically lower than Dice coefficients or correlations due to the impact of changing either map on the total score.

### Testing the Function of the Co-activated Default Mode Network

The function of the co-activated DMN can then be assessed by comparing the time course of the identified DMN component(s) to the task model. The beta coefficients were extracted per condition, based on the SPM design matrix. A one-way ANOVA with condition as the factor was used to assess whether the activity in a specific component related to the task model.

Planned contrasts were implemented if the task model was significant. In each task a critical comparison was used to determine whether the DMN was involved in the condition of interest (*Episodic Task;* episodic>control; *Social - Person Knowledge;* social (visual faces and auditory names)>non-social (visual and auditory animals and landmarks); *Social - Theory of Mind;* social (intentional) >non-social (random) motion). All tasks were plotted against rest. Bonferroni correction was applied for the resulting three planned contrasts each for the *Episodic Task* and the *Social - Theory of Mind Task.* For the *Social - Person Knowledge* data, it would be possible to find no difference between the social and non-social semantic conditions because of involvement of the co-activated DMN in both social and non-social semantics. Therefore, this possibility is assessed by additionally contrasting the social and non-social conditions to a non-semantic baseline. For this dataset Bonferroni correction was applied for six planned contrasts. The Bonferroni-adjusted p-values are reported in all cases.

The main focus of this paper is to test hypothesised functions of the network-of-interest, the DMN. Therefore, the full pattern of results is shown for the network-of-interest. However, these findings may be complemented with an understanding of the networks responsible for episodic and social processing. Therefore, all non-artefactual components were checked to assess whether they significantly related to the task model. Of these task-related models, all those that showed significantly greater activity for the task of interest than the critical baseline condition are presented.

## Results

### Determining the Components in each Task Dataset

Based on the group mean Minimum Description Length criteria estimates, 26 components were identified in the *Episodic Task* dataset, 31 in the *Social - Person Knowledge* dataset and 20 in the *Social - Theory of Mind* dataset. Artefactual components were removed from each dataset (*Episodic Task, 13 artefactual components; Social - Person Knowledge, 17 artefactual components; Social - Theory of Mind, 9 artefactual components).* Components in each dataset are referred to as E (*Episodic Task*), P (*Social - Person Knowledge Task)* or T *(Social - Theory of Mind Task*) followed by their associated number. All non-artefactual components are shown in Supplementary Figures 1, 2 and 3.

### Identifying the Network-of-Interest within each Task Dataset

Each non-artefactual component was quantitatively compared to the DMN templates using the Jaccard similarity coefficient. Comparison to the DMN identified in Jackson et al., (2019) detected a single component above threshold in each task dataset (*Episodic Task, E9, J = .209; Social - Person Knowledge, P18, J =.174; Social - Theory of Mind, T8, J =.235).* In the *Social - Theory of Mind Task* two other components had a Jaccard similarity coefficient greater than the initial threshold of .15 (T13; J=.161, T2; J=.157). However, the combination of the first component (T8) with these additional components reduced the similarity with the template, therefore only T8 was selected as reflecting the DMN. The DMN template provided in Shirer et al., (2012) identified the same components (*Episodic Task, E9, J = .191; Social - Person Knowledge, P18, J =.158; Social - Theory of Mind, T8, J =.207),* confirming successful detection of a DMN consistent with the prior literature. In the *Social - Theory of Mind Task* an additional component reached the initial selection threshold (T13; J=.164), yet combining the two components reduced the similarity with the template, therefore only T8 was selected. Figure 1 shows the overlap between the component identified in each dataset and the Jackson et al., (2019) template. Overlap with the Shirer (2012) template may be seen in Supplementary Figure 4. All identified components include the core DMN regions and their peak activations are listed in Table 1.

**Figure 1.**
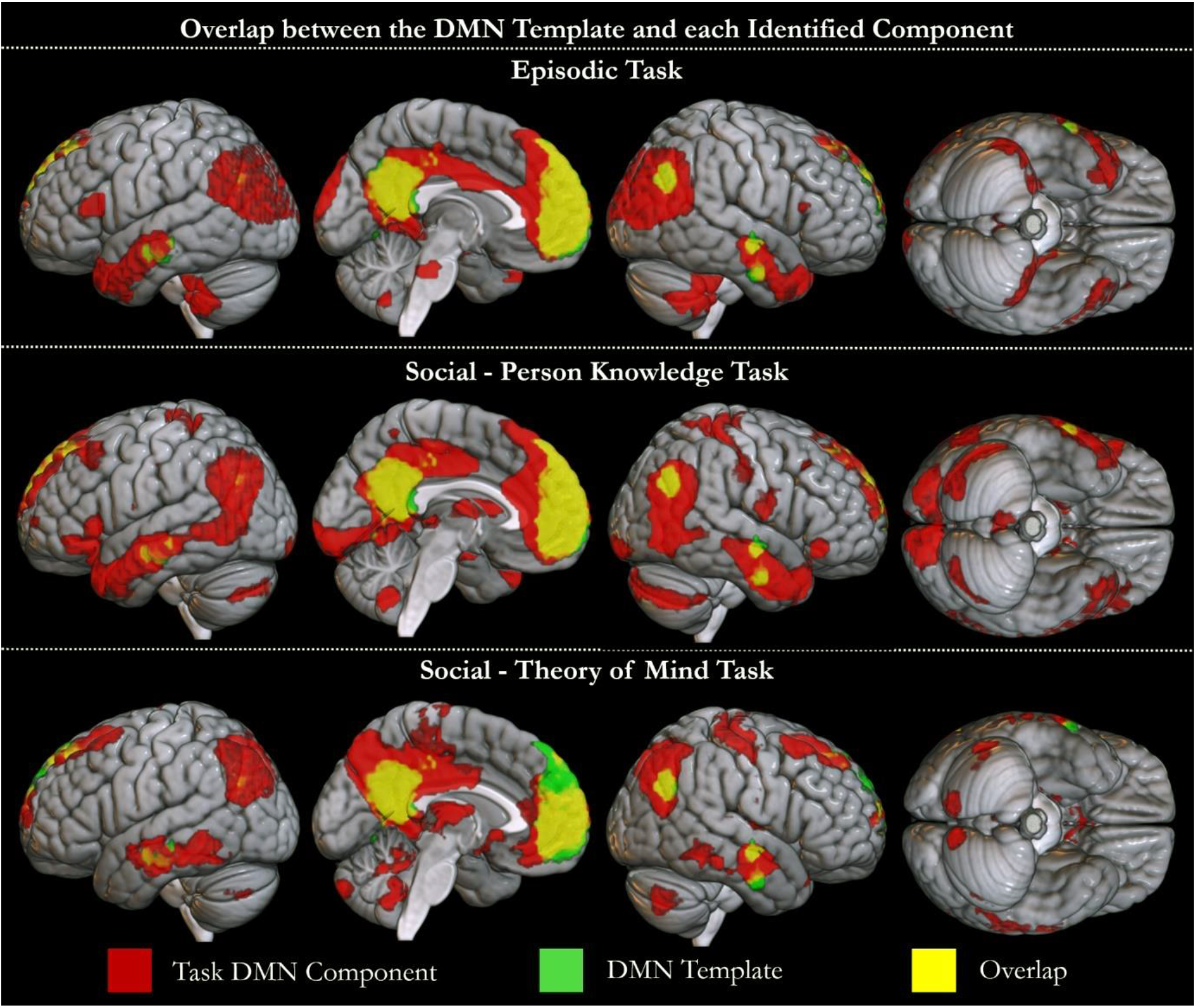
The spatial overlap between the *a priori* template obtained from Jackson et al., (2019) and the component identified to be the DMN in each task is shown. The components identified in the *Episodic Task* (E9), *Social - Person Knowledge Task* (P18) and *Social - Theory of Mind Task* (T8) datasets are shown in red. The template is shown in green. Overlap is shown in yellow.

**Table 1.**
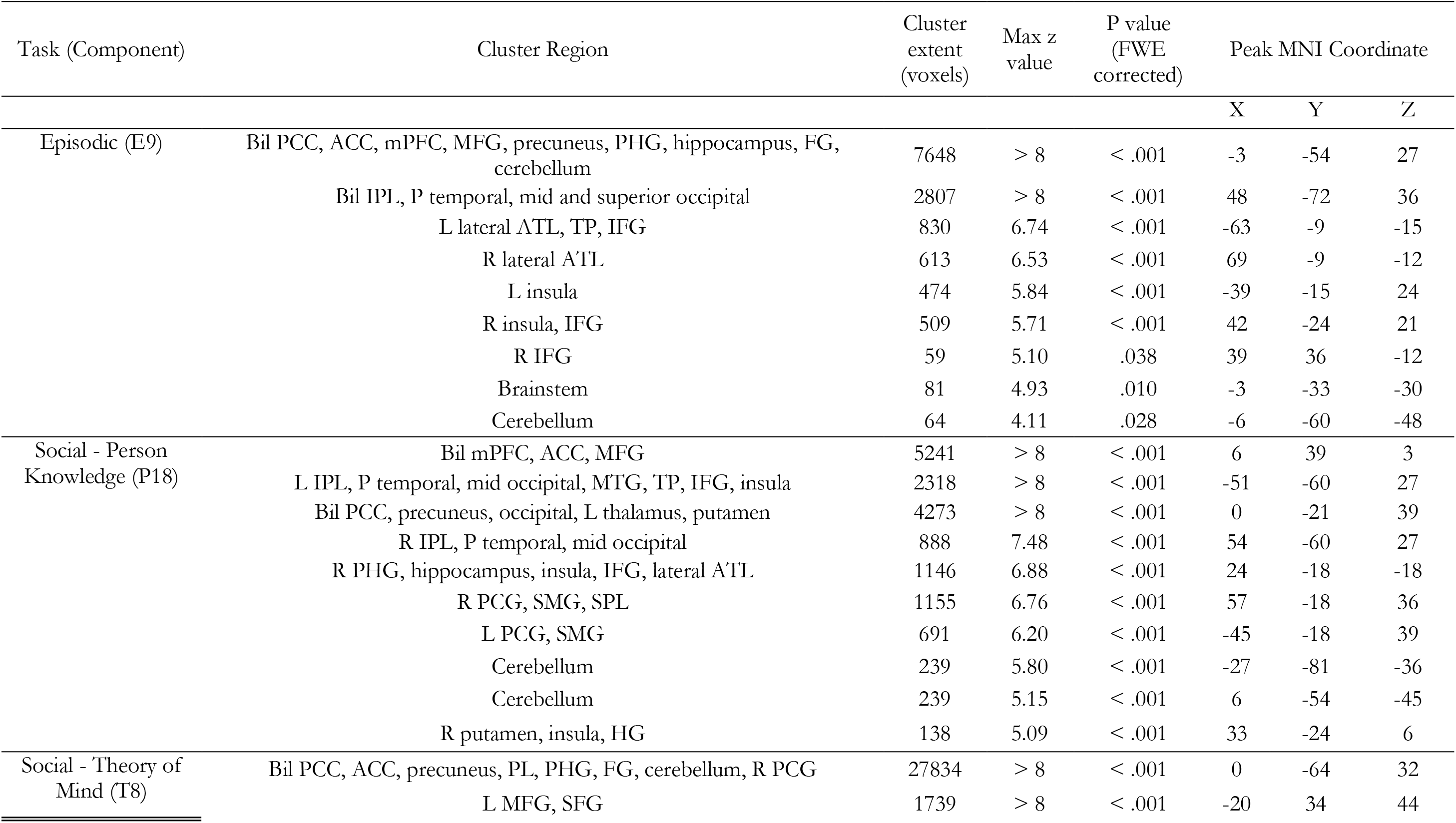
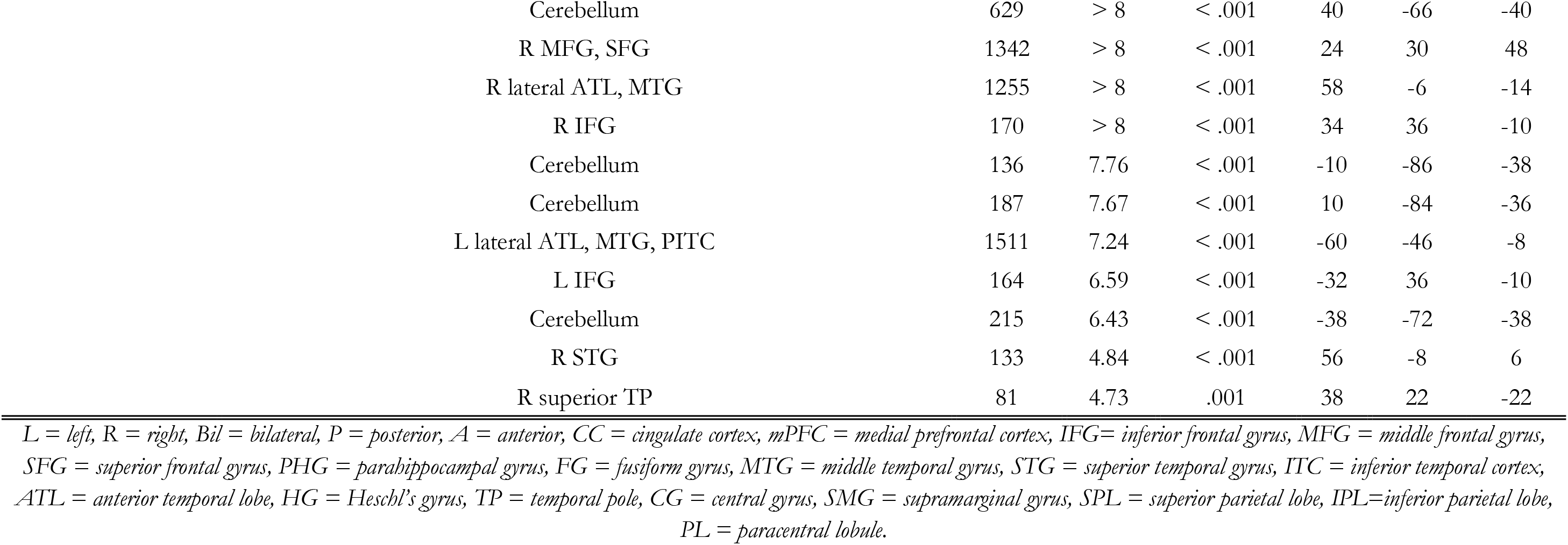
Peak activation in the DMN component-of-interest in each task dataset. Voxels are significant at .001. Clusters are significant using FWE- correction and a critical cluster level of 0.05.

### Testing the Hypothesised Functions of the Co-activated DMN Episodic Task Results

The DMN component (E9) is shown in Figure 2.A. This component was significantly task-related (F(4, 105)=43.138, p<.001). The critical contrast of episodic > control processing showed significantly less activity in the DMN for the episodic than the control condition (t(21) = -7.817, p<.001). This provides no evidence for the involvement of the DMN in explicit episodic retrieval tasks. Both the episodic and control condition were deactivated compared to rest, significantly for the episodic condition (t(21) = -10.249, p<.001) and as a trend from control (t(21) = -2.431, p=.050). The behavioural profile of the DMN component is shown in Figure 2.A.

**Figure 2.**
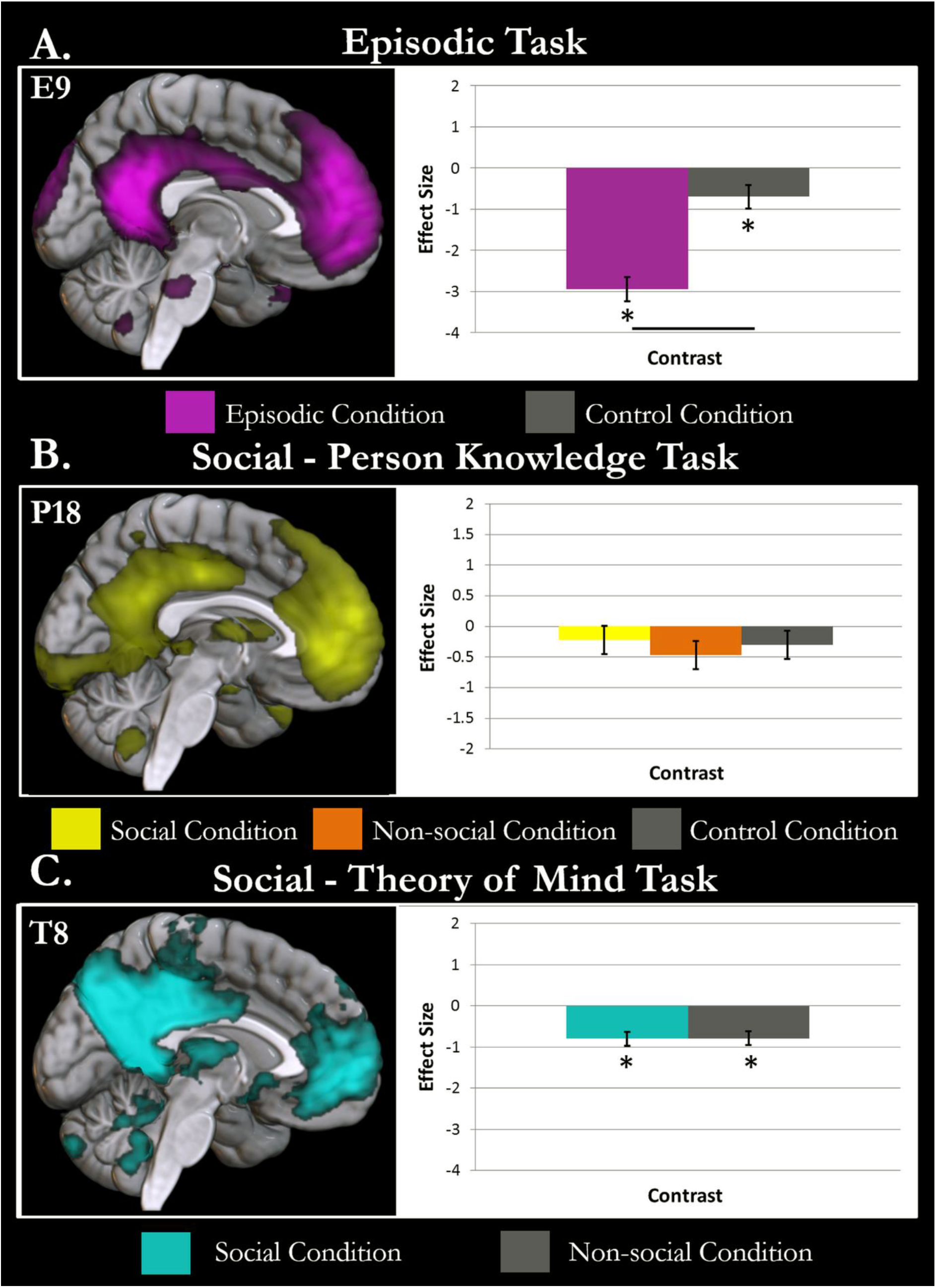
The task relation of the DMN component identified in each task is shown. Conditions with activity significantly different to rest (p<.05 after Bonferroni adjustment) are highlighted with an asterisk. Differences between conditions are demonstrated by a black line between the condition bars. A. The *Episodic Task* DMN component is displayed in violet. The involvement of the component in the episodic (violet) and baseline conditions (grey) is displayed against rest. B. The DMN component identified in the *Social - Person Knowledge Task* is displayed in yellow. The activity of the component in the social semantic (yellow), non-social semantic (orange) and baseline conditions (grey) are shown compared to rest. C. The DMN component identified in the *Social - Theory of Mind Task* is shown in cyan. The activity of this component in the social motion (cyan) and non-social motion conditions (grey) are shown against rest. The co-activated DMN was not found to be significantly more active in the condition of interest (episodic/social processing) than the baseline condition for any task.

To further understand the results of univariate analyses of episodic data, all non-artefactual components with a significant relation to the task model, were assessed for their involvement in episodic processing. The contrast episodic > control processing was significant for components E15, E19, E20 and E25. Each of these components is shown in Figure 3 and the areas of peak activation are listed in Table 2. E15 involved frontal and parietal regions with a left dominance and may reflect the left frontoparietal control network (Vincent et al. 2008). E19 includes a large swathe of dorsomedial prefrontal cortex and anterior cingulate cortex, as well as supramarginal gyrus, frontal and occipital areas. E20 was primarily centred on the precuneus and bilateral angular gyri, the defining features of the precuneus network (Margulies et al. 2009; Uddin et al. 2009). E25 involves activity in occipital, posterior inferior temporal, motor and posterior cingulate cortices, as well as a small region of the medial prefrontal cortex. The full behavioural profile of these components is shown in Figure 3. All were activated for the episodic condition compared to rest (E15; t(21)= 19.497, p<.001; E19; t(21)= 9.778, p<.001; E20; t(21)= 6.017, p<.001; E25; t(21)= 19.655, p<.001) and all except E15 showed significantly greater activation for the control condition than rest (E15; t(21)= -2.258, p=.078; E19; t(21)= 4.663, p<.001; E20; t(21)= 3.128, p<.001; E25; t(21)= 16.913, p<.01). This may reflect some (albeit lesser) involvement of these components in the visual task. Overall, these results suggest a key role for multiple networks in episodic cognition, including the precuneus network and left frontoparietal networks. These networks involve much of the medial surface, however, this is the result of distinct episodic components involving the posterior and anterior DMN regions. There is no evidence the co-activated DMN is involved in episodic retrieval. However, many of the areas in or near the DMN do appear to be involved in episodic memory as part of other (sometimes multiple) networks.

**Figure 3.**
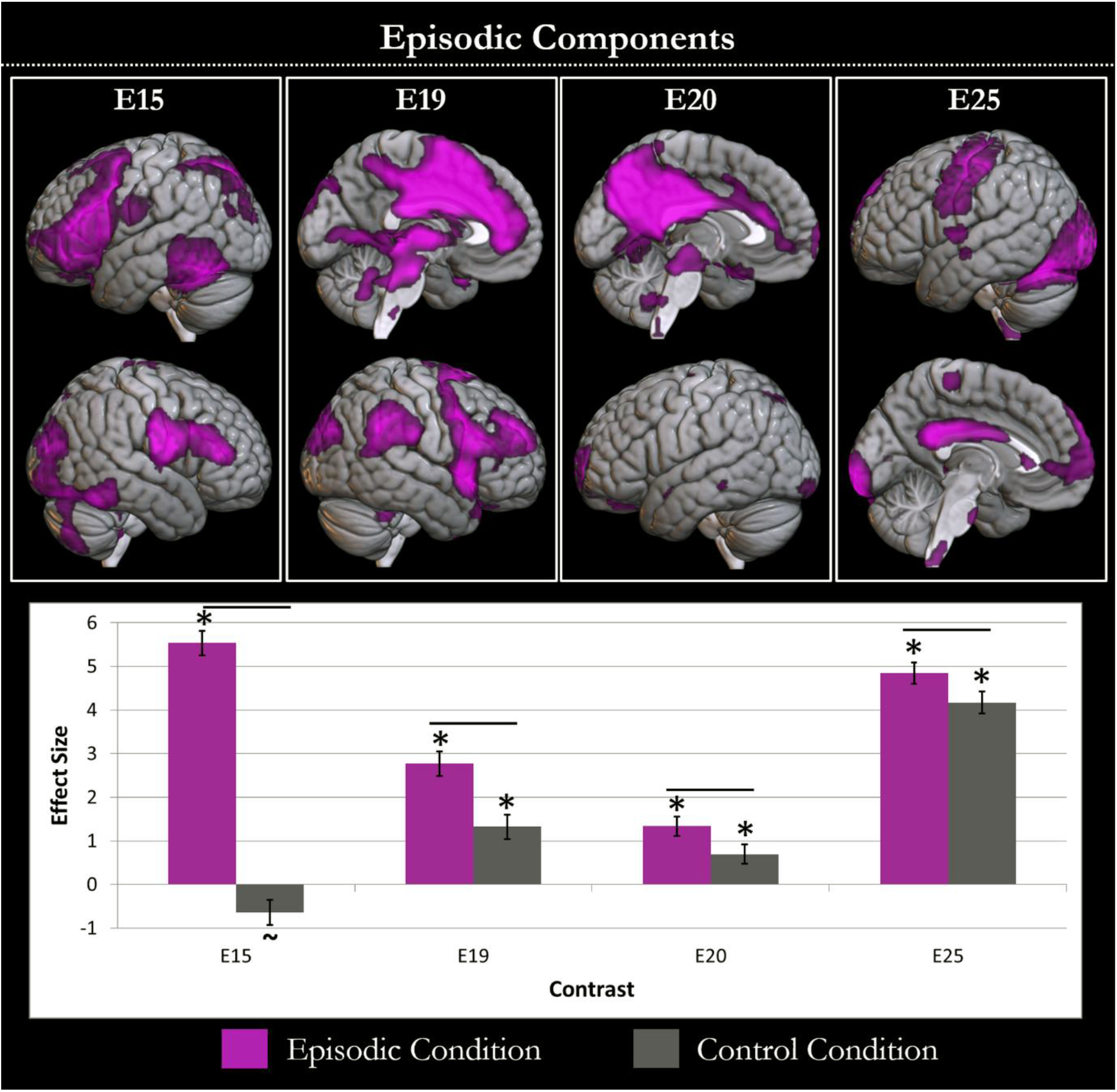
The components with greater involvement in the episodic than the baseline condition are displayed. Each component is shown in violet in the top panel and the task relation of the component is shown in the lower graph. The graph shows the effect size of each component in the episodic (violet) and baseline (grey) conditions compared to rest. Conditions with activity significantly different (p<.05 after Bonferroni adjustment) to rest are highlighted with an asterisk. A tilde is shown if there is a trend (p<.1 after Bonferroni adjustment) towards this difference. Differences between conditions are demonstrated by a black line between the condition bars.

**Table 2.**
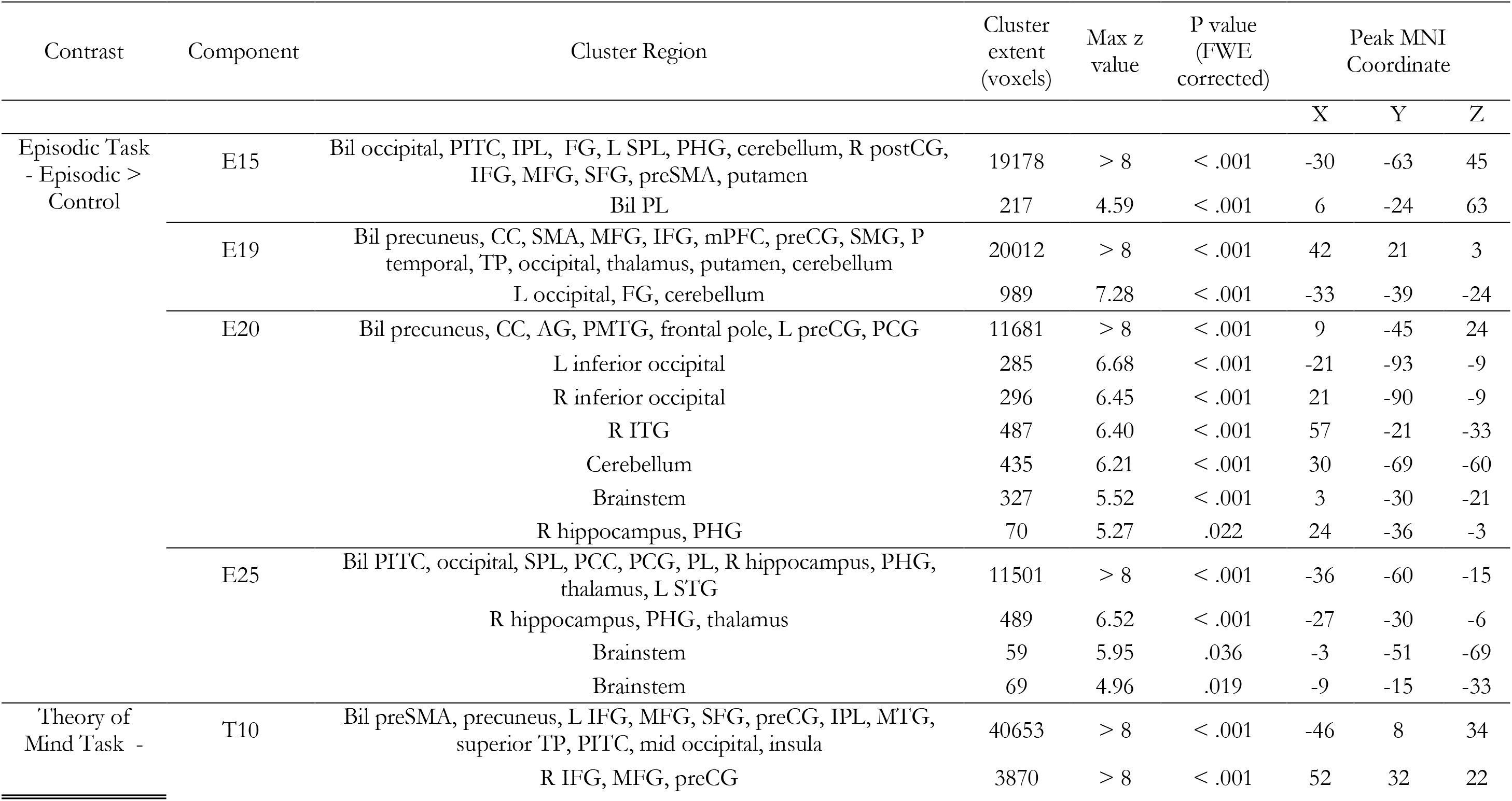
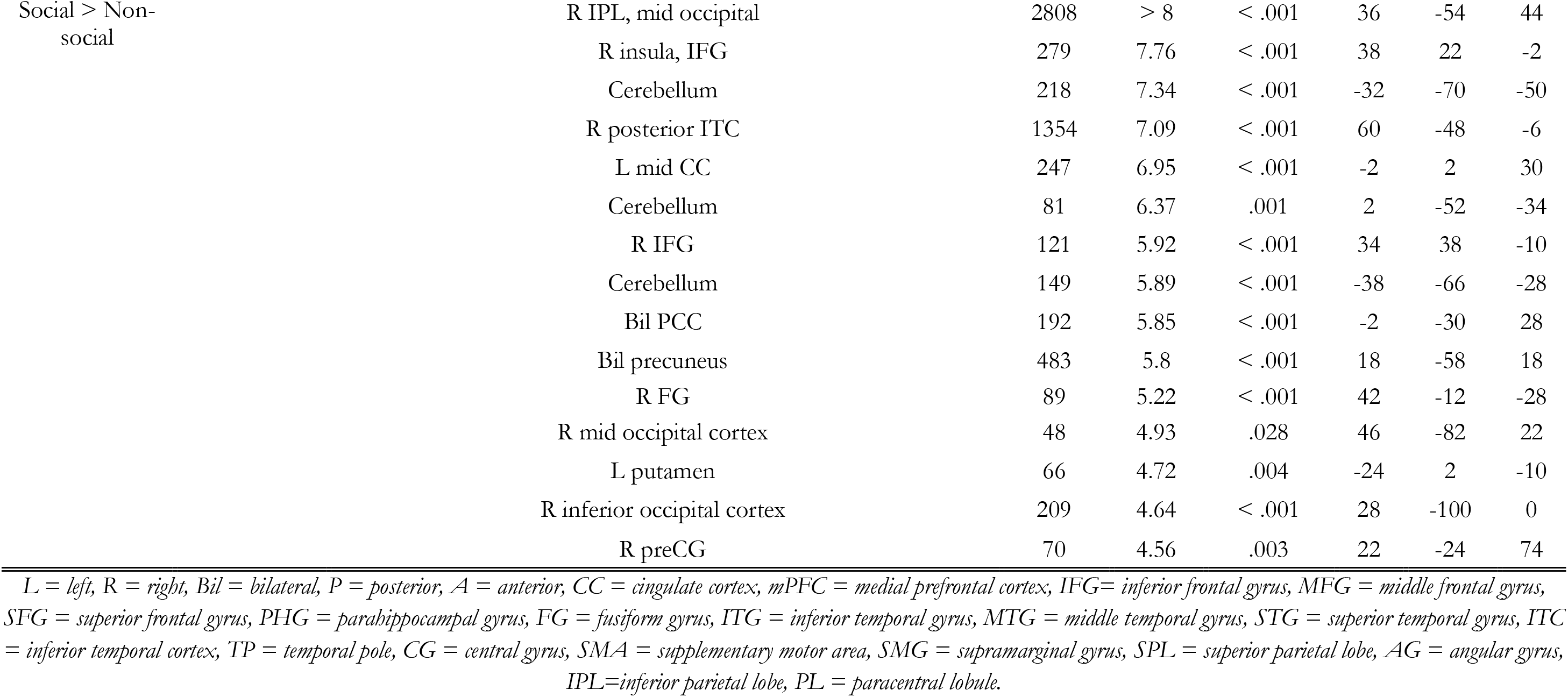
Peak activation in the components identified with each contrast-of-interest. Voxels are significant at .001. Clusters are significant using FWE-correction and a critical cluster level of 0.05.

### Social - Person Knowledge Task Results

Figure 2.B. shows the DMN component identified in the *Social - Person Knowledge Task* data (P18). The DMN was not significantly task-related (F(3,68)=1.425, p=.243) therefore the results of the contrasts should not be interpreted. However, for completeness in assessing the core goal of the current endeavour the contrasts are reported here with caution. The contrast social > non-social was not significant (t(18)=1.074, p=1). These results provide no evidence that the DMN is specifically involved in the kind of social cognition necessary to perform person knowledge judgements. Neither the social or non-social conditions were significantly different to the non-semantic control condition (social>control; t(18) = 0.364, p=1; non-social>control; t(18) = - 0.711, p=1). There was no evidence the co-activated DMN was involved in either social or general semantics, providing no evidence that the DMN is necessary for social tasks of this nature. In addition, this independent dataset shows no involvement of the co-activated DMN in semantic cognition, supporting the conclusions from Humphreys et al., (2015) and directly replicating the findings of Jackson et al., (2019). Compared to rest, both the social (t(18) = - 0.954, p=1) and the non-social condition (t(18) = -2.028, p=.279) show non-significant deactivation of the co-activated DMN.

No other components were significantly more active in the social condition compared to the non-social condition, even without correction for multiple comparisons. This does not imply that there were no components involved in the social task, but merely that none were involved to a significantly greater extent when the relevant knowledge was related to people. Indeed, the same components were recruited equally for the social and non-social semantic conditions (P12; F(3,68) = 8.253, p<.001, social *vs.* control t(18)=3.974, p<.005, non-social *vs*. control t(18)= 2.817, p<.05, social *vs*. non-social t(18)=1.157, p=1; P31; F(3,68) = 17.713, p<.001, social *vs*. control t(18)=2.978, p<.05, non-social *vs*. control t(18)= 3.415, p<.01, social *vs*. non-social t(18)=-0.437, p=1). P12 involved bilateral temporal cortices and inferior frontal gyri, areas critical for general semantic cognition. P31 included occipital and motor cortex and may reflect the processing of visual input and motor output. Both are shown in Supplementary Figure 2.

### Social - Theory of Mind Task Results

The DMN component (T8) identified in the *Social - Theory of Mind Task* is displayed in Figure 2.C. alongside the activity of the social and non-social conditions (plotted against rest). The DMN was significantly related to the task model (F(2,237)=15.297, p<.001). The contrast social > non-social was not significant (t(79)=-0.047, p=1). Therefore, no evidence was provided for the hypothesis that the DMN is involved in theory of mind. The DMN was deactivated compared to rest for viewing motion with or without social interaction (social>rest; t(79)=-4.814, p<.001; non-social>rest; t(79)=-4.766, p<.001).

All non-artefactual components were assessed for their relation to theory of mind processing. One component showed a significant effect for the social > non-social contrast (T10; F(2, 237)=48.393, p<.001; t(79)=3.448, p<.005). T10 was significantly activated for both social (t(79)=9.704, p<.001) and non-social (t(79)=6.256, p<.001) motion over rest. The spatial extent and behavioural profile of T10 is shown in Figure 4. The locations of peak activations in T10 are recorded in Table 2. T10 is left hemisphere-biased and includes bilateral inferior and middle frontal gyri, dorsomedial prefrontal cortex, inferior parietal lobe and posterior inferior temporal cortex. All regions had a greater extent in the left hemisphere and there was additional involvement of left middle and superior temporal gyri. This component appears to reflect the left frontoparietal control network and is involved in processing both kinds of motion, yet to a much greater extent when processing the social condition. As there is no high-level baseline in this dataset it is not possible to accurately assess the networks involved in both social and non-social conditions (although others may well be involved). Therefore, no further results are presented.

**Figure 4.**
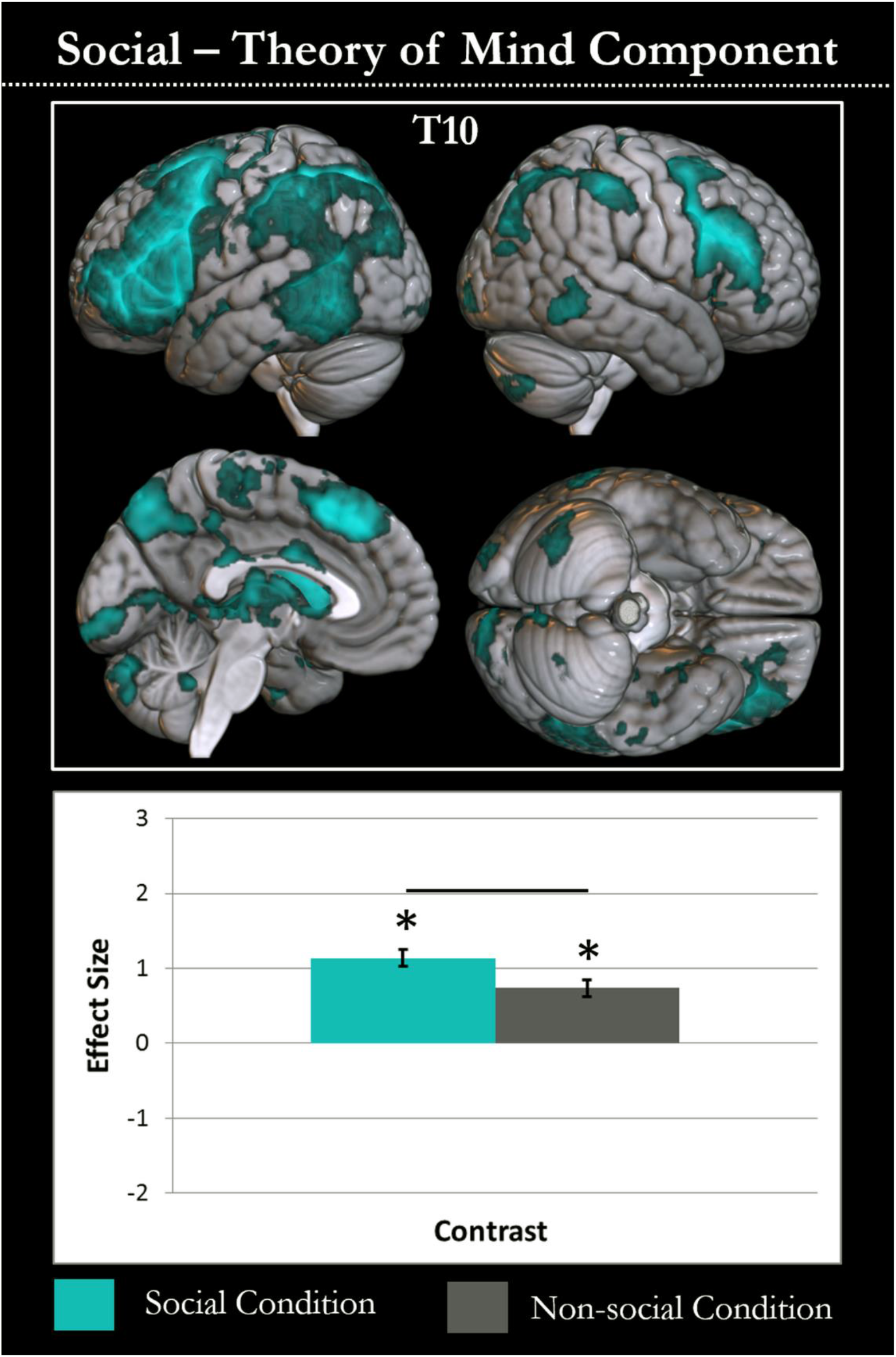
The single component with greater involvement in the theory of mind than the baseline condition is displayed. The spatial extent of the component is shown in cyan in the top panel and the task relation of the component is shown in the lower graph. The graph shows the effect size of each component in the social motion (cyan) and non-social motion conditions (grey) compared to rest. Conditions with activity significantly different (p<.05 after Bonferroni adjustment) to rest are highlighted with an asterisk. Differences between conditions are demonstrated by a black line between the condition bars.

## Discussion

Applying a network-level approach to episodic and social task data, we consistently observed a co-activated set of regions that were positively identified as the co-activated DMN, but found no evidence for involvement of this network in episodic or social cognition. Indeed, this co-activated DMN showed the reverse of the pattern one would expect from an episodic network, being significantly less active in episodic than visual baseline tasks, and all tasks showed less DMN involvement than rest. Instead these tasks relied on distinct sets of regions, including other known RSNs. Indeed, both episodic memory and theory of mind recruited a left-biased frontoparietal network (FPN), linked to executive control processing, planning and information load and typically associated with a task-positive activation pattern (Vincent et al. 2008; Spreng et al. 2010b; Summerfield et al. 2010; Meyer et al. 2012; Spreng 2012). Multiple networks contributed to episodic processing, perhaps supporting different processes. Social semantics relied on the same network as non-social semantic cognition. In the remainder of the Discussion, we consider the co-activated networks found to relate to episodic and social cognition, why prior studies have implicated the DMN, and the possible consequences for its functional role.

### What co-activated networks support episodic and social processing?

Visual assessment suggests the co-activated sets of regions supporting episodic memory include the precuneus network, consisting of dorsal precuneus and bilateral angular gyri (Damoiseaux et al. 2006; Buckner et al. 2008; Margulies et al. 2009; Stevens et al. 2009; Zhang et al. 2012; Yang et al. 2013; Utevsky et al. 2014), and the left FPN, as well as additional components focussed on frontal, occipital, cingulate, inferior temporal and motor regions. Whilst traditional episodic memory research focussed on the MTL, these results are highly compatible with the increasing move toward recognising the involvement of multiple distributed areas or networks, including precuneus, angular gyrus and frontal cortex (Buckner et al. 2001; Rugg 2002; Lundstrom et al. 2005; Wagner et al. 2005; Kim 2010; Ranganath et al. 2012; Hirose et al. 2013; Jeong et al. 2015). Person-knowledge relied on the same semantic network as knowledge of non-social concepts, which corresponded extremely well to prior region and network-level assessments of semantic cognition (Binder et al. 2011; Jackson et al. 2016; Lambon Ralph et al. 2017), including the lack of involvement of the co-activated DMN (Jackson et al. 2019). Although univariate comparison may find subtle regional differences between social and non-social judgements, these are in the wider context of similar networks and may relate to difficulty level (Zahn et al. 2007; Binney et al. 2016; Rice et al. 2018; Binney et al. 2020). In contrast, theory of mind processing was only associated with differences in the left FPN. This FPN component includes the temporoparietal and dorsomedial prefrontal areas highlighted as critical for mentalising in prior univariate analyses, including analyses of this task (Castelli et al. 2000; Castelli et al. 2002; Wheatley et al. 2007; Barch et al. 2013). However, FPN involvement may relate to task difficulty differences and may not be the only network responsible for completion of this task (as intentional and non-intentional motion could recruit additional shared networks).

### Why do DMN regions appear in region-level episodic and social analyses?

The co-activated DMN was not found to be involved in the episodic or social tasks, despite previously being linked to these domains on the basis of region-level and partial least squares analyses (e.g., Buckner et al. 2008; Kim 2010; Spreng et al. 2010a; Spreng et al. 2010b; Mars et al. 2012; Hassabis et al. 2014; DuPre et al. 2016; DiNicola et al. 2020). There are multiple possible reasons why activation differences could be identified in relevant areas without involvement of the co-activated DMN, and distinct factors may be critical for the different tasks. Many of the constituent regions of the DMN are involved in episodic memory as part of distinct networks, such as the FPN and the precuneus network. Therefore, activation in DMN regions for episodic memory may relate to the function of one (or, if overlapping, more than one) of these networks. The default mode and precuneus networks have been separated by ICA (Damoiseaux et al. 2006; Stevens et al. 2009; Zuo et al. 2010) and distinguished on the basis of function (Utevsky et al. 2014), including the relation of their constituent regions to episodic memory (Sestieri et al. 2011). Here we support this distinction by demonstrating that the precuneus network, but not the co-activated DMN supports episodic memory. Both the precuneus and the angular gyri are commonly associated with episodic processing (Lundstrom et al. 2003; Lundstrom et al. 2005; Wagner et al. 2005; Cavanna et al. 2006; Cabeza et al. 2008; Dorfel et al. 2009; Humphreys et al. 2014), yet whether this was due to the activity of the core DMN or precuneus network was not previously clear. The presence of multiple partially overlapping or nearby networks in many DMN regions could have led the co-activated DMN to be incorrectly associated with episodic memory, due to the importance of one or more of these distinct networks for episodic memory.

Difficulty differences could lead to relatively greater time spent in free thought at the end of trials, or a reduced need to suppress off-task thoughts associated with the DMN (Gilbert et al. 2012; Humphreys et al. 2017). This may be a critical factor resulting in the identification of activity differences in DMN regions for social versus non-social semantics. Relative differences could be induced within the DMN via the comparison of easier with harder items or tasks, particularly if these are consistent across studies (e.g., if processing intentional motion is typically easier than random motion or if comprehending human intentions is easier than assessing physical causality; Spreng 2012; Humphreys et al. 2017). Alternatively, the critical issue may be the accurate identification of the extent of the DMN, including its separation from the nearby semantic network. Meta-analyses of theory of mind have shown involvement of DMN regions (particularly medial prefrontal cortex), however the focus may be outside of core DMN regions (e.g., temporoparietal junction; Buckner et al. 2008; Van Overwalle 2009; Mars et al. 2012) and explicit overlap is minimal (Mars et al. 2012). The current findings are in high accord with research associating episodic and social processing with distinct sets of regions (referred to as DMN ‘subnetworks’) and not the full DMN, including the resemblance of the areas implicated in social processing to the semantic network, and the identification of regions typically linked to both DMN and FPN for episodic processing (Andrews-Hanna et al. 2014; DiNicola et al. 2020).

### What implications are there for the functional role of the regions that preferentially activate during rest?

It is important to consider how these assessments of the co-activated DMN relate to the wider set of regions showing greater activation in rest than many tasks (regardless of connectivity or co-activation). As well as the co-activated DMN, this pattern may be found for multiple networks identified here including the precuneus and semantic networks, possibly due to the high propensity for episodic and semantic cognition during rest (Binder et al. 1999; Buckner et al. 2008; Binder et al. 2009; Humphreys et al. 2015). Each of these networks were separated based on their connectivity and had distinct cognitive profiles. This is highly compatible with a componential view of these task-negative regions, suggesting they do not reflect one coherent network, but multiple component parts with dissociable functions, such as episodic, social and semantic cognition. This idea is supported by a great deal of prior research using a range of methods to demonstrate that there are multiple different task-negative networks with distinct relations to function (e.g., Andrews-Hanna et al. 2010; Leech et al. 2011; Spreng 2012; Humphreys et al. 2015; Axelrod et al. 2017; DiNicola et al. 2020). Unlike parcellation methods employed to directly ‘split’ the DMN (e.g., Andrews-Hanna et al. 2010), the networks implicated here are not restricted to include only areas that fall within a particular definition of the DMN and a single DMN region can be included in multiple networks across time.

The identification of multiple networks that include task-negative regions, is often conceptualised as splitting the DMN into distinct subnetworks. Thus, it may be tempting to conclude that the broad task-negative DMN includes subnetworks responsible for episodic, social and semantic processing, albeit with the function of at least one subnetwork (the core co-activated DMN) unknown (e.g., Andrews-Hanna et al. 2010). However, some aspects of the current and prior research may urge caution in applying a strong version of this particular interpretation. Firstly, the regions which typically deactivate during tasks are ill-defined as the precise set of regions that deactivates varies based on the task or tasks used, which could result in radically different ‘default mode networks’ (Callard et al. 2012; Humphreys et al. 2015).

Indeed, even primary sensory regions deactivate when other senses are engaged (e.g., Humphreys et al. 2015). Secondly, there is not strong functional connectivity between all of these regions (this is why they can be separated). Indeed, they may only appear functionally connected due to a single element of their functional response; their high activity in rest. Thirdly, splitting the task-negative regions into ‘subnetworks’ responsible for different functions ignores the identification of additional regions that do not typically deactivate during tasks in these co-activated networks. These factors make definition and interpretation of a broad multi-component DMN difficult, resulting in a lack of clarity on how to determine which networks should be considered subnetworks of the DMN and what implications this has for their interpretation. It is possible that these proposed subnetworks may have little in common, outside of including some regions which deactivate during some tasks. In addition, it should be noted that despite the possibility of splitting the DMN into different ICA components, a set of co-activated regions that included the core DMN regions could be quantitatively identified as matching a priori DMN templates in all datasets. Thus, the presence of these ‘subnetworks’ did not lead to the loss of a core co-activated DMN by splitting it apart. Therefore, whilst these issues may encourage us to be cautious in associating the full set of task-negative regions with a particular function or even set of functions, we can ask; what function does the co-activated DMN support?

### What is the function of the co-activated DMN?

If the co-activated DMN is not engaged in episodic, social or semantic cognition what is its functional role? It is possible that this core DMN forms a distinct ICA component without being a co-activated network with a function, purely as it is a useful way to explain a lot of variance in the datasets. However, this seems unlikely due to the strong structural and functional connectivity between the core midline regions (which is not explained by the other networks identified) and the extremely high level of consistency in finding the DMN in different task and rest datasets (e.g., Greicius et al. 2009). One possibility is the DMN simply performs some specific episodic or social process not tested here, for instance, episodic memory encoding, self-referential thought or processing highly emotional stimuli. An important caveat of the current work if that only one episodic task probing recall was used and the involvement of different networks may shift between differentiable episodic subprocesses (Hirose et al. 2013; Jeong et al. 2015). Further testing at the network-level may find evidence supporting a more specific role in episodic memory, or the co-activated DMN may be responsible for processing in a different domain that we have yet to consider. Alternatively, it may be that this core DMN uses information provided by the semantic and episodic networks, yet their combination is more than the sum of its parts. For instance, distinct combinatorial processes may be required to produce a train of thought (Humphries et al. 2007; Graves et al. 2010; Price et al. 2015) or additional mental simulation may be needed to imagine the future using past events (Buckner et al. 2007; Binder et al. 2011). The core DMN may work in concert with related networks to use episodic and semantic information in a complex manner (Irish et al. 2013) and support ongoing thought more generally (Smallwood et al. 2021). Until this puzzle is resolved, satisfactory interpretations of relevant research, including the implications of clinical differences in the DMN, will not be possible. Further work should investigate different possible functional roles of the co-activated DMN using a network-level approach.

## Acknowledgements

Data were provided [in part] by the Human Connectome Project, WU-Minn Consortium (Principal Investigators: David Van Essen and Kamil Ugurbil; 1U54MH091657) funded by the 16 NIH Institutes and Centers that support the NIH Blueprint for Neuroscience Research; and by the McDonnell Center for Systems Neuroscience at Washington University.

## Data and code availability

code used for this study is available from the following public repository github.com/JacksonBecky/ICA. The data is held in a repository by the MRC Cognition & Brain Sciences Unit and available upon request.

