## Supplementary Materials for "A Network-level Test of the Role of the Co-activated Default Mode Network in Episodic Recall and Social Cognition"

### The Coherent Default Mode Network is not involved in Episodic Memory or Social Cognition

**Supplementary Materials**

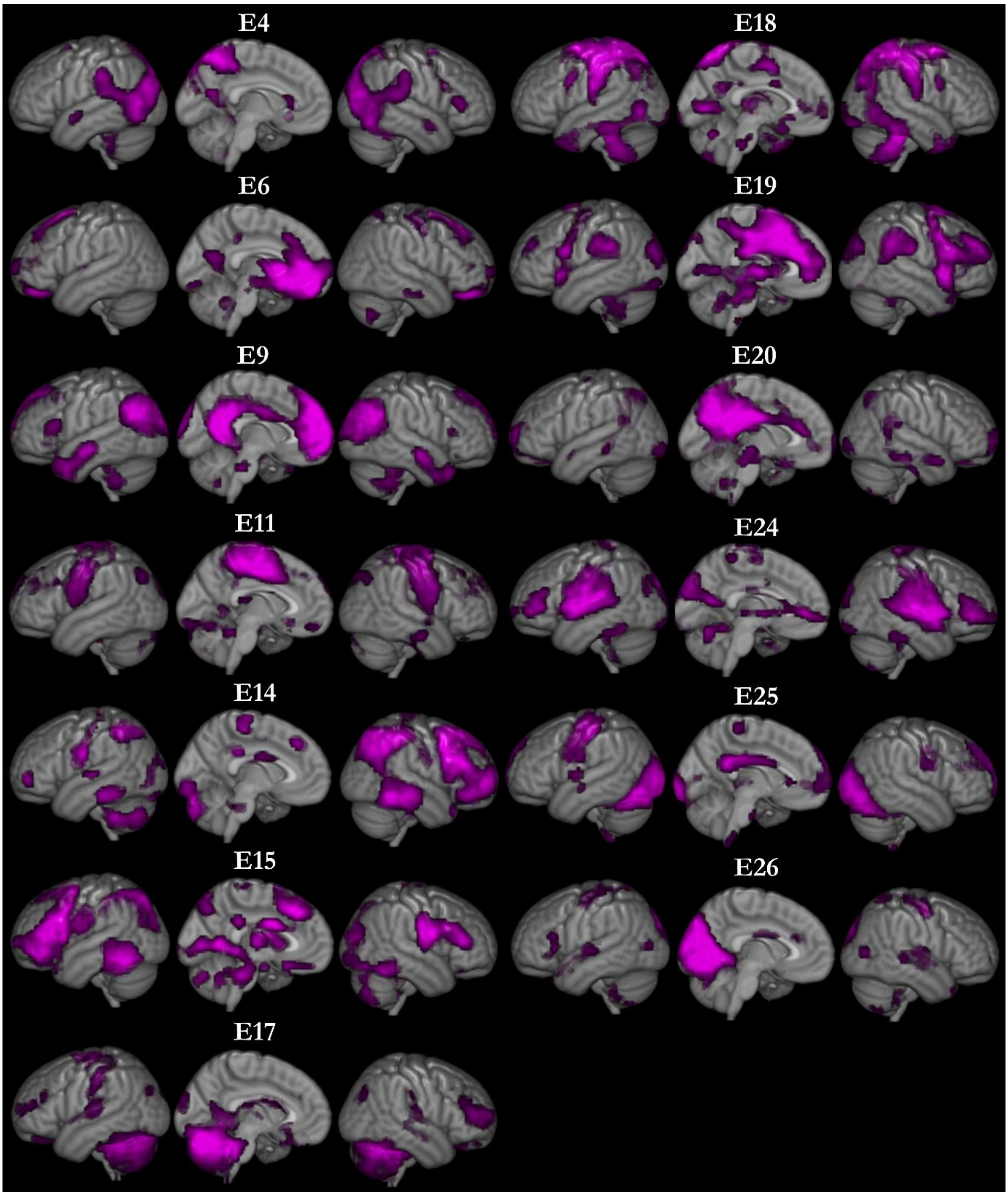


*Supplementary Figure 1*. Each non-artefactual component identified in the *Episodic Task* dataset is shown in violet with its corresponding label.

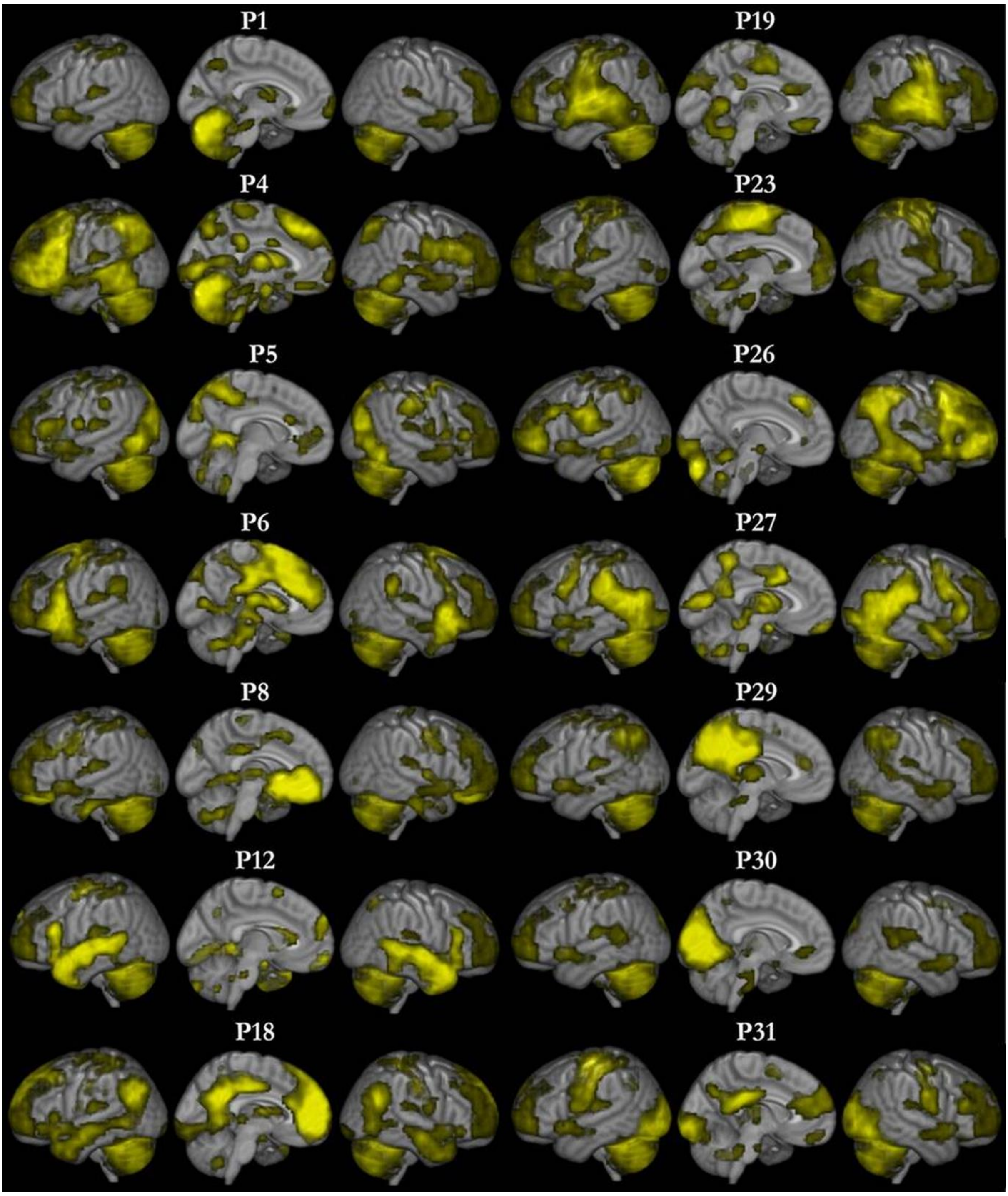


*Supplementary Figure 2*. Each non-artefactual component identified in the *Social - Person Knowledge Task* dataset is shown in yellow with its corresponding label.

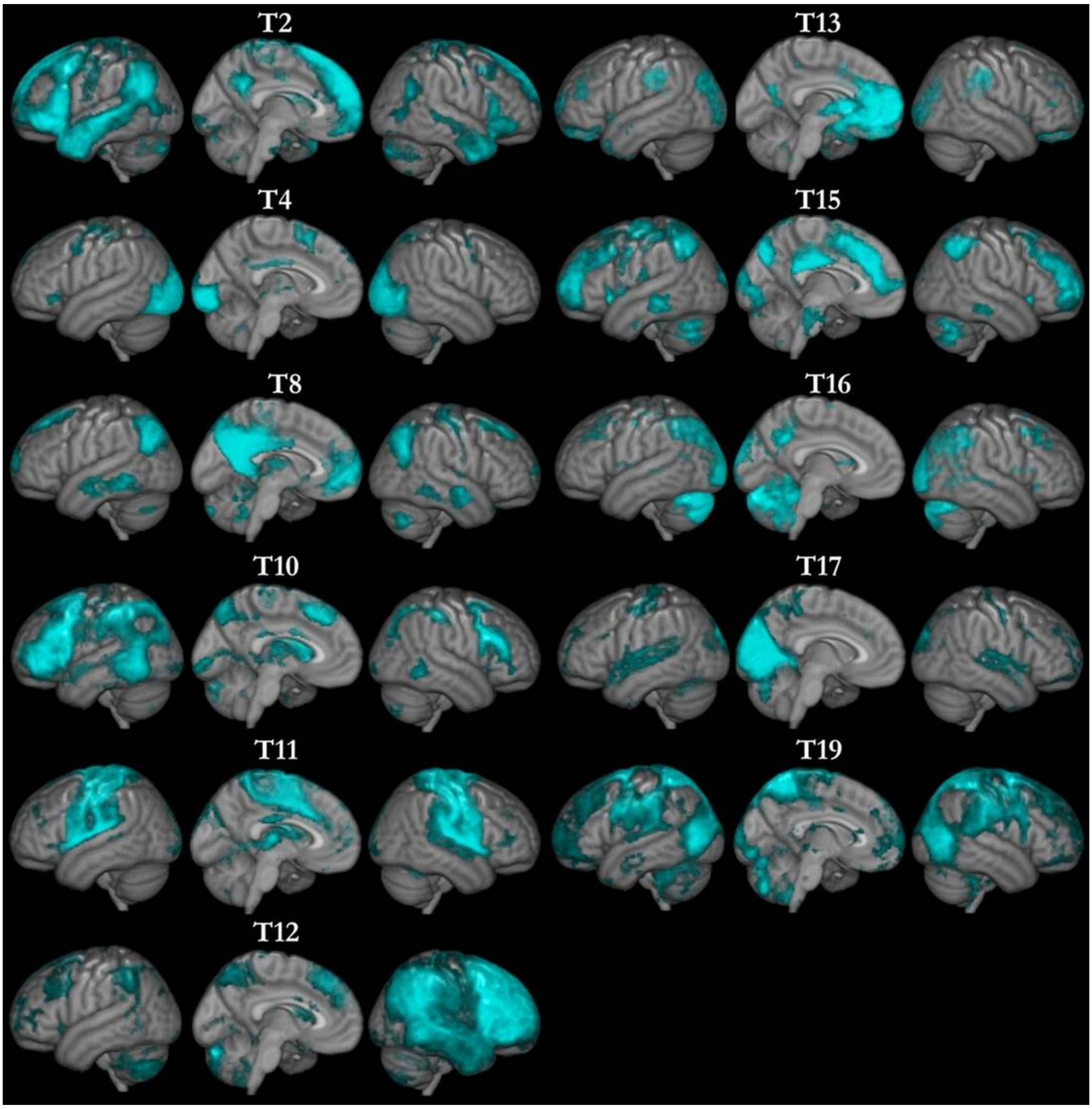


*Supplementary Figure 3*. Each non-artefactual component identified in the *Social - Theory of Mind Task* dataset is shown in cyan with its corresponding label.

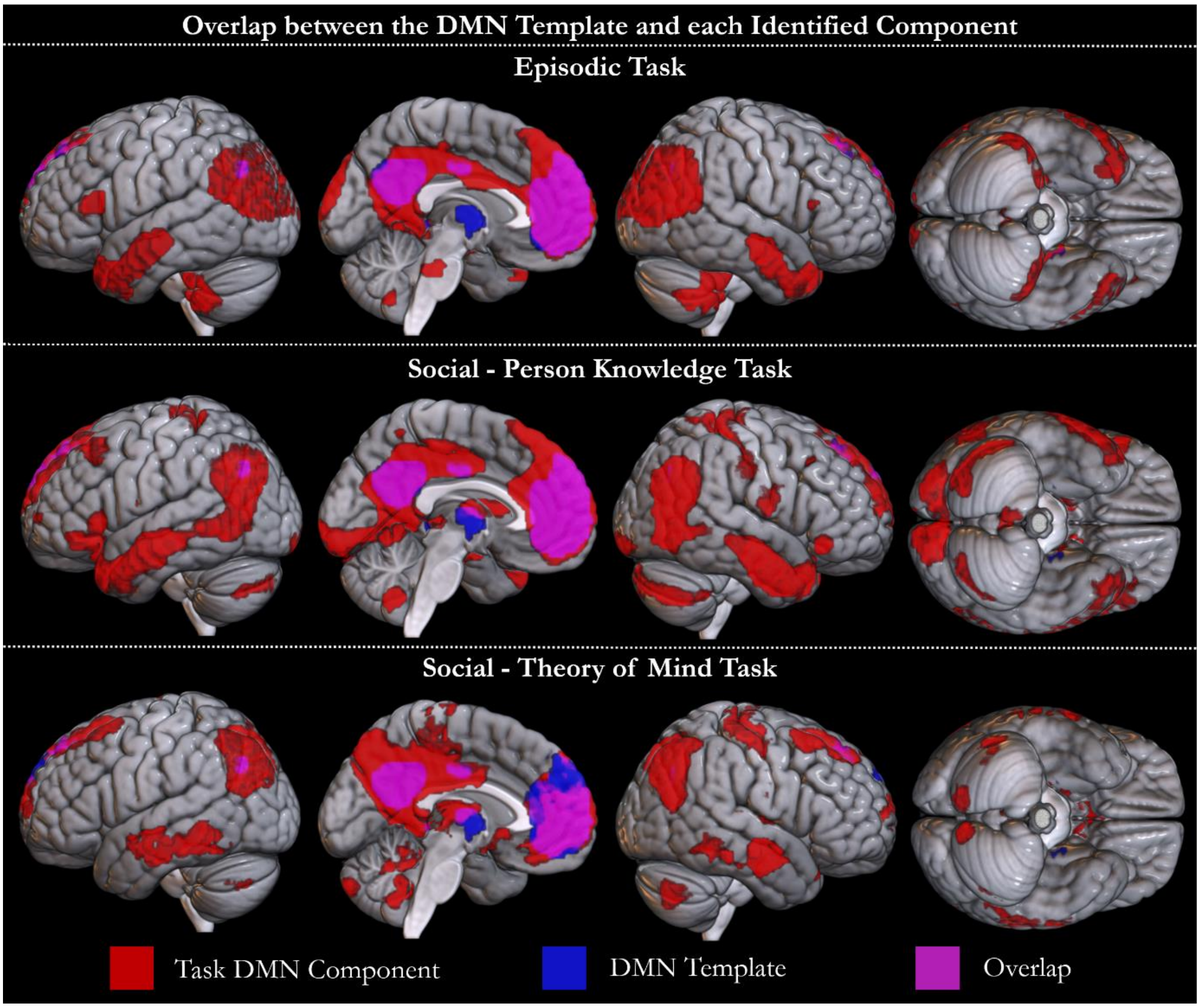


*Supplementary Figure 4.* The spatial overlap between the *a priori* template obtained from Shirer et al., (2012) and the component identified to be the DMN in each task is shown. The components identified in the *Episodic Task* (E9), *Social - Person Knowledge Task* (P18) and *Social - Theory of Mind Task* (T8) are shown in red. The template is shown in blue. Overlap is shown in violet.
